# SCRIPT: predicting single-cell long-range *cis*-regulation based on pretrained graph attention networks

**DOI:** 10.1101/2025.04.27.650894

**Authors:** Yu Zhang, Baole Wen, Yifeng Jiao, Yuchen Liu, Xin Guo, Yushuai Wu, Jiyang Li, Limei Han, Yinghui Xu, Xin Gao, Yuan Qi, Yuan Cheng, Ying He, Weidong Tian

**Affiliations:** Artificial Intelligence Innovation and Incubation Institute, Fudan University, Shanghai, China; Shanghai Academy of Artificial Intelligence for Science, Shanghai, China; INF Technology (Shanghai) Co. Ltd, Shanghai, China; State Key Laboratory of Genetics and Development of Complex Phenotypes, Department of Computational Biology, School of Life Sciences, Fudan University, Shanghai, China; Computer, Electrical and Mathematical Sciences and Engineering Division, King Abdullah University of Science and Technology (KAUST), Thuwal, Saudi Arabia; Center of Excellence for Smart Health, King Abdullah University of Science and Technology (KAUST); Thuwal, Saudi Arabia; Center of Excellence on GenAI, King Abdullah University of Science and Technology (KAUST); Thuwal, Saudi Arabia; Zhongshan Hospital, Fudan University, Shanghai, China; Children’s Hospital of Fudan University, Shanghai, China; Children’s Hospital of Shandong University, Jinan, China

## Abstract

Single-cell *cis*-regulatory relationships (CRRs) are essential for deciphering transcriptional regulation and understanding the pathogenic mechanisms of disease-associated non-coding variants. Existing computational methods struggle to accurately predict single-cell CRRs due to inadequately integrating causal biological principles and large-scale single-cell data. Here, we present SCRIPT (Single-cell *Cis*-regulatory Relationship Identifier based on Pre-Trained graph attention networks) that infers single-cell CRRs from transcriptomic and chromatin accessibility data. SCRIPT incorporates two key innovations: graph causal attention networks supported by empirical CRR evidence, and representation learning enhanced through pretraining on atlas-scale single-cell chromatin accessibility data. Validation using cell-type-specific chromatin contact data demonstrates that SCRIPT achieves a mean AUC of 0.9, significantly outperforming state-of-the-art methods (AUC: 0.68). Notably, SCRIPT obtains a threefold improvement in predicting long-range CRRs (>100 Kb) compared to existing methods. Applying SCRIPT to Alzheimer’s disease and schizophrenia, we establish a framework for prioritizing disease-causing variants and elucidating their functional effects in a cell-type-specific manner. By uncovering molecular genetic mechanisms undetected by existing computational methods, SCRIPT provides a roadmap for advancing genetic diagnosis and target discovery.

## Introduction

The human genome harbors an extensive repertoire of *cis*-regulatory elements (CREs) which are preferentially located in accessible chromatin regions, orchestrating precise spatial and temporal patterns of gene expression^1^. A major mechanism by which CREs, such as enhancers, act on a target gene is through chromosomal looping, bringing distal enhancers into close physical proximity with their target genes within three-dimensional (3D) chromatin space^2^. Notably, the *cis*-regulatory relationships (CRRs) between CREs and their corresponding target genes exhibit two defining characteristics: (1) CRRs are highly cell-type-specific, reflecting the unique transcriptional programs of distinct cellular contexts^3^; and (2) for more than half of CRRs, the genomic distances between CREs and their target genes exceed 100 kilobases (Kb)^4^. Consequently, single-cell-resolution CRR data, particularly those capturing long-range CRRs, is essential for understanding the regulatory mechanism underlying cell-type-specific gene expression^3, 5^. Moreover, genome-wide association studies (GWASs) have identified numerous variants associated with complex diseases^6^, the majority of which are located in non-coding regions^7^. Single-cell long-range CRR data can link more non-coding variants to their target genes in a cell-type-specific manner, providing deeper insights into the genetic underpinnings of complex diseases^5, 8^.

Extensive CRR information can be obtained through high-throughput chromosome conformation capture (Hi-C) technique^9^ and expression quantitative trait loci (eQTLs) analysis^10^. However, these datasets are predominantly available at the tissue level rather than the single-cell level. To address this limitation, single-cell Hi-C (scHi-C) technologies have been developed to generate single-cell CRR data^11^. However, their high cost and the limited availability of publicly accessible scHi-C datasets hinder widespread adoption of them.

An alternative approach involves developing tools to predict single-cell CRRs by leveraging the growing abundance of single-cell assay for transposase-accessible chromatin using sequencing (scATAC-seq) and single-cell RNA sequencing (scRNA-seq) data. Current tools, such as SCARLink^12^ and LINGER^13^, still face critical challenges. First, these tools use correlation-based algorithms to infer single-cell CRRs without considering the casual biological principles, which might contribute to their insufficient accuracy in identifying long-range CRRs. Second, they utilize a limited amount of single-cell data for model training rather than exploit the wealth of large-scale publicly available single-cell atlas datasets. The limited training data are not enough to learn complex single-cell *cis-*regulatory mechanisms.

To address these challenges, we propose a novel deep-learning-based method called SCRIPT (Single-cell *Cis-*regulatory Relationship Identifier based on Pre-Trained graph attention networks). SCRIPT incorporates two key algorithmic innovations. First, it leverages graph causal attention networks (GCATs) to simulate *cis-* transcriptional regulation, using chromatin accessibility of CREs measured by scATAC-seq to predict gene expression levels quantified by scRNA-seq via CRRs. Causal attention masks of GCATs^14, 15^ are designed by incorporating empirical evidence from large-scale bulk Hi-C and eQTL datasets, enabling the modeling of *cis*-transcriptional regulation grounded in biological principles. Second, SCRIPT employs a self-supervised graph autoencoder (SSGAE)^16^ pretrained on atlas-scale scATAC-seq data to comprehend the complex interactions between CREs across diverse tissues. This pretraining enables the model to generate effective CRE representations. Recent advancements in foundation models for single-cell transcriptomics prove the effectiveness of the pretraining strategy^17, 18^. Relying on these innovative algorithm designs, SCRIPT exhibits excellent performance in predicting single-cell CRRs, as validated by cell-type-specific chromatin contact data. Compared to current computational methods, SCRIPT identifies disease-causing variants more accurately and explains their functional effects more reasonably at the cell type level. Furthermore, we provide a SCRIPT-based framework to unravel the pathogenic mechanisms underlying the non-coding variants associated with complex diseases, including Alzheimer’s disease (AD) and schizophrenia (SCZ). SCRIPT holds promise in advancing genetic diagnosis and therapeutic target discovery for various complex genetic diseases.

## Result

### The overview of SCRIPT

The biological basis of *cis*-regulation is chromosomal looping, wherein distal CREs are brought into close physical proximity with their target genes within 3D space. The core concept of SCRIPT is incorporating the biological principles of *cis*-regulation into graph neural networks to simulate *cis*-regulatory mechanisms. The biological principles are from comprehensive empirical CRR evidence composed of large-scale tissue-level Hi-C (from 27 human tissues) and eQTL (from 49 human tissues) datasets (Fig. 1a).

**Fig. 1.**
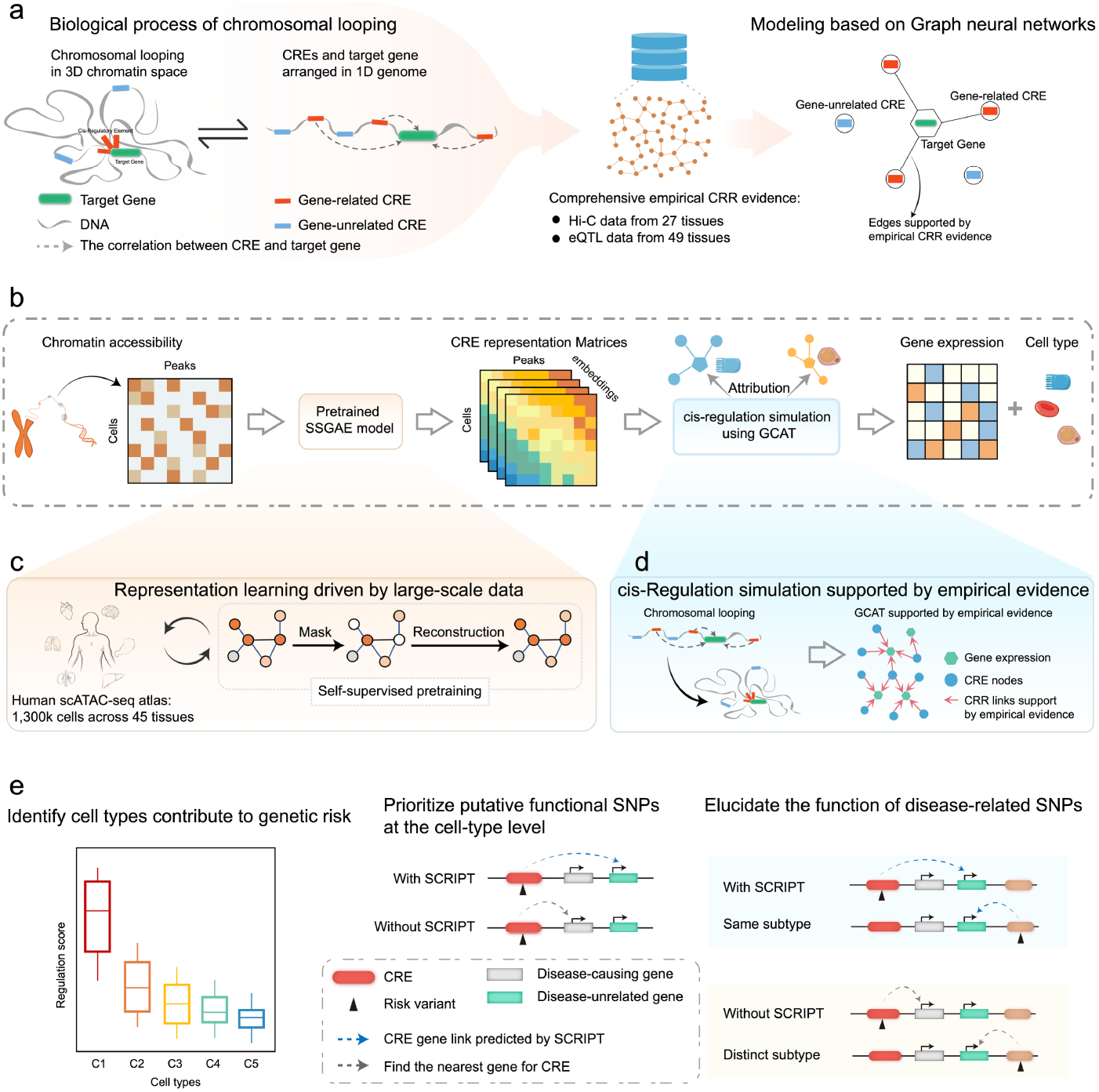
Overview of SCRIPT. **a,** Graph neural networks inspired by biological principles. SCRIPT incorporates biological principles into graph neural networks to simulate *cis*-regulatory mechanisms. The biological principles are from comprehensive empirical CRR evidence composed of large-scale Hi-C and eQTL datasets. **b,** Schematic illustration of SCRIPT. SCRIPT uses chromatin accessibility data as input to predict gene expression and cell type, leveraging the pretrained SSGAE and GCAT modules. Single-cell regulation scores are generated from the GCAT module using the attribution method. **c,** Representation learning driven by large-scale data. The SSGAE is pretrained on atlas-scale scATAC-seq datasets, enabling the generation of biologically meaningful CRE representations. **d,** *Cis*-regulation simulation supported by empirical evidence. GCAT is designed to mimic chromosomal looping to utilize the CRE representations to predict gene expression. **e,** Applications in disease biology. SCRIPT facilitates the identification of cell types contributing to genetic risk, prioritization of candidate disease-causing variants, and elucidation of the function of disease-related variants. CRR, *cis-*regulatory relationship; CRE, *cis-*regulatory element; Hi-C, high-throughput chromosome conformation capture; eQTL, expression quantitative trait loci; SSGAE; self-supervised graph autoencoder; GCAT, graph causal attention network; scATAC-seq, single-cell assay for transposase-accessible chromatin using sequencing.

The *cis*-regulation simulation of SCRIPT leverages chromatin accessibility data to predict gene expression and cell type. Regulation scores for single-cell CRRs are inferred through an attribution method applied to the trained simulation model, which assesses the impact of each CRE-gene link on its corresponding gene. SCRIPT’s accurate predictions are driven by two innovative components: the pretrained SSGAE model and GCAT supported by empirical evidence (Fig. 1b; for details, see Methods and Fig. S1).

SSGAE is pretrained on atlas-scale human scATAC-seq data (∼1.3 million cells), enabling it to generate biologically meaningful representations for previously unseen scATAC-seq datasets (Fig. S2). During pretraining, SSGAE randomly masks node features in the input data and encodes the corrupted graph into node embeddings using an encoder. A decoder then reconstructs the masked input node features. By pretraining on large-scale scATAC-seq data, the SSGAE gains a fundamental understanding of *cis*-regulatory networks, encoding the network hierarchy into the model’s attention weights in a completely self-supervised manner (Fig. 1c).

The algorithmic design of GCAT mimics chromosomal looping to utilize the CRE representations to predict gene expression. During GCAT training, attention scores of edges representing CRE-gene links, supported by empirical evidence, are updated directionally from CREs to target genes. GCAT enables SCRIPT to predict single-cell CRRs based on biological principles, rather than simple data correlations (Fig. 1d).

SCRIPT provides a new avenue for exploring the pathogenic mechanisms of complex diseases. Its applications include the identification of cell types contributing to genetic risk, prioritization of candidate disease-causing variants, and elucidation of the function of disease-related variants (Fig. 1e).

### SCRIPT outperforms existing methods in cell-type-specific CRR prediction

To evaluate SCRIPT’s performance, cell-type-specific chromatin contact data (proximity ligation-assisted chromatin immunoprecipitation sequencing (PLAC-seq)^19^ and cell-type-specific promoter capture Hi-C (pcHi-C)^20^) are regarded as the true labels, and four pairs of scATAC-seq and scRNA-seq datasets derived from brain and blood tissues, not appearing in pretrain datasets, are curated for cell-type specific CRR predictions. Two metrics are employed for comparative analysis across different computational methods: cell-level area under the receiver operating characteristic curve (AUC), and reg-level AUC. The cell-level AUC measures the ability to accurately identify cell-type-specific CRRs, whereas the reg-level AUC evaluates the accuracy of assigning a cell-type-specific CRR to the correct cell type (for details, see Methods and Fig. S3).

To investigate the effectiveness of pretraining in CRR prediction, we conduct a comparative analysis between SCRIPT and SCRIPT without pretraining. The results indicate that pretraining significantly enhances CRR prediction performance, with mean cell-level AUC increasing from 0.85 to 0.9 and mean reg-level AUC rising from 0.69 to 0.74. (Fig. 2a-c, Fig. S4).

**Fig. 2.**
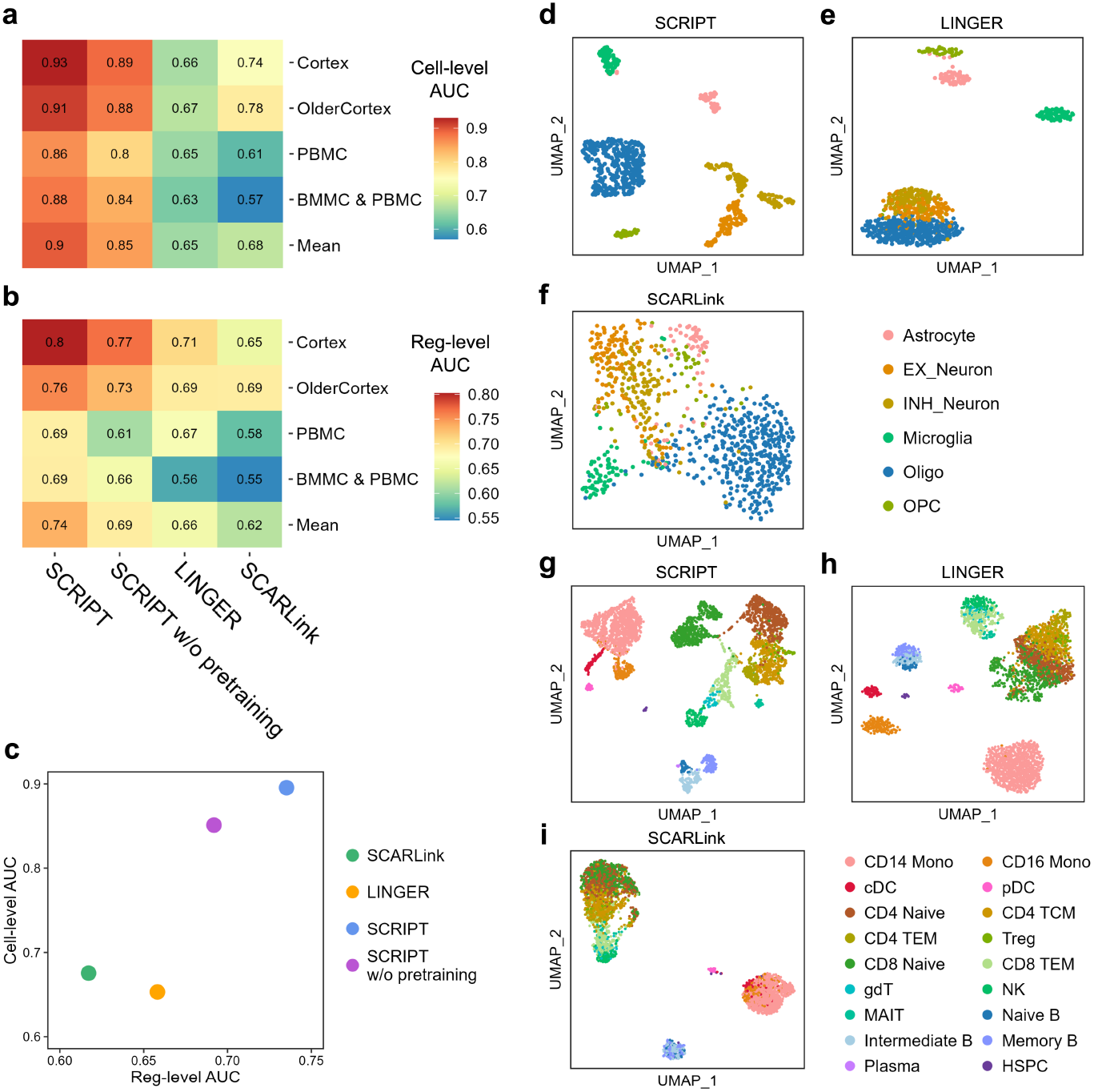
Performance comparison of SCRIPT and competing methods for CRR prediction. **a, b,** Heatmaps showing cell-level **(a)** and reg-level **(b)** AUCs of four methods for single-cell CRR prediction across four benchmark datasets. Method names and dataset names are shown at the bottom and left of the heatmaps, respectively. **c,** Scatter plot of mean AUCs for the four methods, where the *x*-axis represents the mean reg-level AUC and the *y*-axis represents mean cell-level AUC. Ideal outcomes would lie in the top right corner. **d, e, f,** UMAP visualization of single-cell regulation score matrices predicted by SCRIPT **(d)**, LINGER **(e)**, and SCARLink **(f)** for the Cortex dataset (*n* = 852 cells). **g, h, i,** UMAP visualization of single-cell regulation score matrices predicted by SCRIPT **(g)**, LINGER **(h)**, and SCARLink **(i)** for the PBMC dataset (*n* = 3,609 cells). PBMC, peripheral blood mononuclear cells; BMMC, human bone marrow mononuclear cells.

Besides, we compare SCRIPT with two state-of-the-art methods, LINGER and SCARLink. SCRIPT achieves an average cell-level AUC of 0.90 and an average reg-level AUC of 0.74. In comparison, LINGER and SCARLink attain average cell-level AUCs of 0.65 and 0.68, respectively, and reg-level AUCs of 0.66 and 0.62, respectively (Fig. 2a-c, Fig. S5-6). These results demonstrate that SCRIPT significantly outperforms both LINGER and SCARLink in both cell-type-specific CRR prediction accuracy and robustness across diverse cell types.

Subsequently, single-cell regulation score matrices generated by the three methods are visualized using Uniform Manifold Approximation and Projection (UMAP) to assess their ability to characterize biological variations inherent in single-cell data. UMAP visualizations in the cortex dataset reveal that SCRIPT separates distinct cell types, whereas LINGER fails to distinguish excitatory neurons from inhibitory neurons, and SCARLink produces the least coherent cell-type groupings (Fig. 2d-f). Similar results are observed in the peripheral blood mononuclear cells (PBMC) dataset (Fig. 2g-i). Overall, SCRIPT outperforms both LINGER and SCARLink in its ability to capture meaningful biological signals.

### SCRIPT demonstrates superior long-range CRR prediction

To evaluate performance on long-range CRR prediction, CRRs are stratified into groups based on the distances between CREs and the transcription start sites (TSSs) of their corresponding target genes. For the CRR groups 0-10Kb, 10-50Kb and 50-100Kb in the cortex dataset, SCRIPT achieves cell-level AUCs ranging from 0.89 to 0.95 and reg-level AUCs ranging from 0.82 to 0.86, which are higher than LINGER (cell-level AUC: 0.75-0.79, reg-level AUC: 0.77-0.81) and SCARLink (cell-level AUC: 0.79-0.82, reg-level AUC: 0.69-0.77) (Fig. 3a, b). Notably, SCRIPT maintains robust performance in long-range CRR groups (100–300 Kb and 0.3–1 Mb), achieving a mean cell-level AUC of 0.90 and a mean reg-level AUC of 0.75. In contrast, LINGER’s mean cell-level AUC significantly drops to 0.5 and mean reg-level AUC to 0.62, while SCARLink’s mean cell-level AUC significantly declines to 0.62 and mean reg-level AUC to 0.58 (Fig. 3a, b). Notably, in the long-range CRR prediction, SCRIPT achieves over a threefold relative increase in cell-level AUC and a twofold relative increase in reg-level AUC compared to LINGER and SCARLink, considering that a random classifier yields an AUC of 0.5. SCRIPT’s superiority is further evident in the PBMC dataset, where it attains a mean cell-level AUC of 0.85 for long-range CRR groups (100–300 Kb and 0.3–1 Mb), significantly outperforming LINGER (0.60) with over a threefold relative improvement (Fig. 3a, b). These findings underscore SCRIPT’s superior capacity to accurately identify and characterize long-range CRRs.

**Fig. 3.**
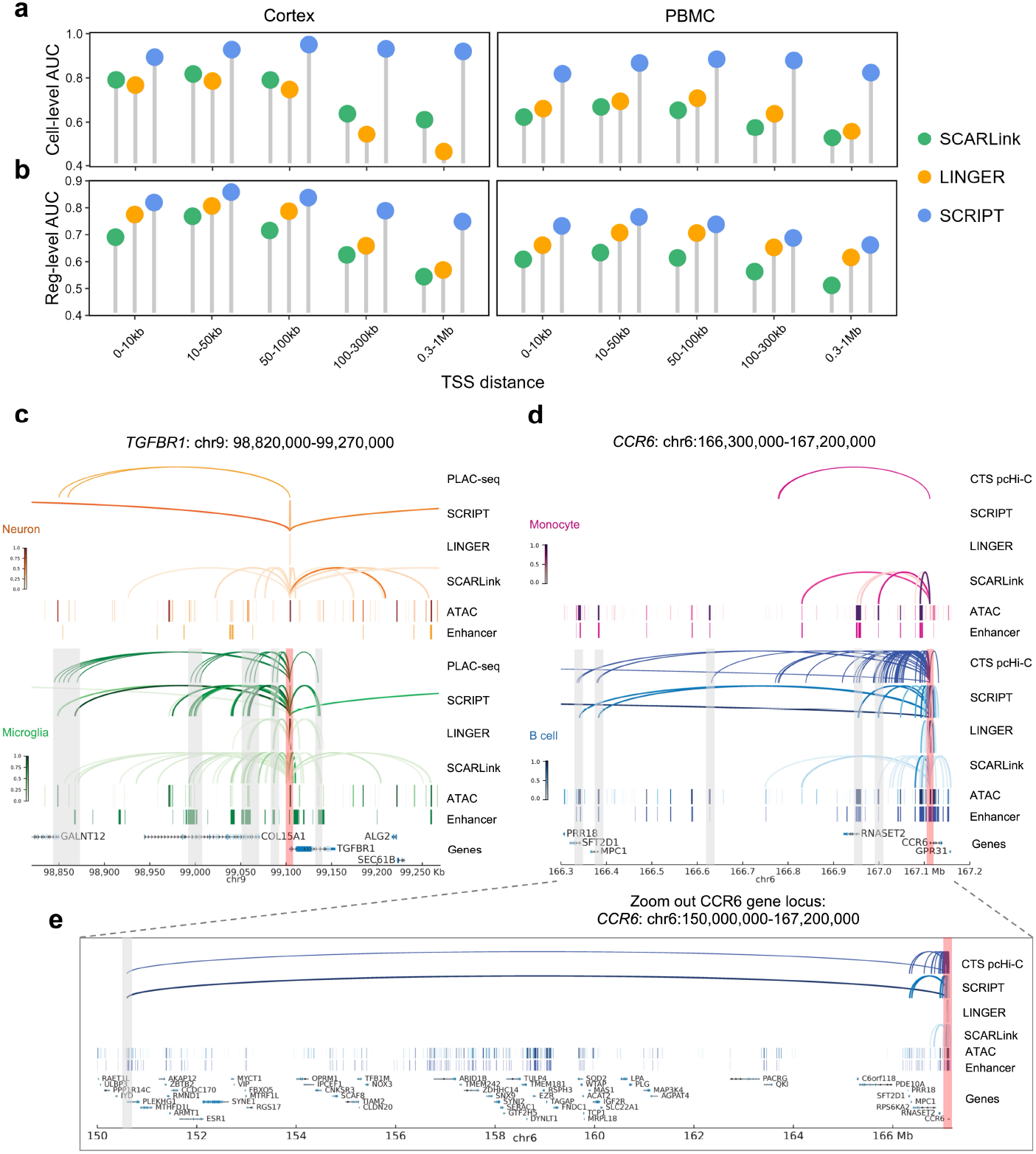
SCRIPT outperforms LINGER and SCARLink in long-range CRR prediction. **a, b** Lollipop plots showing the cell-level **(a)** and reg-level **(b)** AUCs for cell-type-specific CRR prediction in the cortex and PBMC datasets. CRRs are categorized into five distance groups ranging from 0-10Kb to 0.3-1Mb. **c, d,** Genomic visualizations of CRRs identified by PLAC-seq data, normalized regulation scores predicted by SCRIPT, LINGER, and SCARLink, normalized scATAC-seq-derived pseudobulk tracks, and enhancers identified by H3K27ac ChIP-seq data in the *TGFBR1* gene locus (chr9: 98,820,000-99,270,000) **(c)** and *CCR6* gene locus (chr6:166,300,000-167,200,000) **(d)**. The scATAC–seq tracks represent the aggregate chromatin accessibility of all cells from the given cell type and have been normalized for comparability. The CREs linked to *TGFBR1* or *CCR6* based on ChIP-seq and either PLAC-seq or cell-type-specific pcHi-C are highlighted in grey bars. The TSSs of *TGFBR1* and *CCR6* are marked by red bars. **e,** Genomic visualization of the *CCR6* gene locus within a broader genomic context (chr6:150,000,000–167,200,000). CTS pcHi-C, cell-type-specific promoter capture Hi-C.

To further inspect the superiority of SCRIPT to LINGER and SCARLink in long-range CRR prediction, we visualized CRRs identified by PLAC-seq and compared them with predictions made by the three methods. As a case study, we examine the *TGFBR1* locus, a microglia marker gene critical for microglial maturation^21^. In microglia, PLAC-seq data identify five sets of CREs regulating *TGFBR1^19^*, located approximately 250kb upstream, 105kb upstream, 45kb upstream, 15kb upstream, and 30kb downstream from the TSS of *TGFBR1* (highlighted by the grey bars in Fig. 3c). SCRIPT accurately identifies all five CRE sets (Fig. 3c, Table S1), whereas LINGER detects only the three CRE sets closest to the TSS (Fig. 3c). Although SCARLink identifies all five, it produces numerous false-positive predictions. Particularly, SCARLink predicts many CRRs in neurons that are not corroborated by PLAC-seq data, highlighting its reduced specificity compared to SCRIPT (Fig. 3c). Moreover, H3K27ac is a histone modification associated with the higher activation of transcription and therefore considered as an enhancer mark^22^. The CREs identified by SCRIPT to regulate *TGFBR1* are consistently labeled as microglia-specific enhancers in H3K27ac ChIP–seq data^19^. In contrast, many CREs identified by SCARLink fail to align with H3K27ac ChIP–seq data, further emphasizing its tendency to generate false positives (Fig. 3c, Fig. S7a).

Another representative example displays SCRIPT’s advantages in longer-range CRR prediction. *CCR6*, a well-established marker gene for B cells, plays an important role in the cell fate transition of B cells^23^. Cell-type-specific pcHi-C and H3K27ac ChIP– seq data^20, 24^ identify five remote candidate enhancers regulating *CCR6*, located approximately 775Kb, 730Kb, 480Kb, 165Kb, and 125Kb upstream of the TSS of *CCR6* (highlighted by the grey bars in Fig. 3d). SCRIPT successfully detects four of these enhancers (Fig. 3d, Table S2), whereas SCARLink identifies only two, and LINGER fails to identify any (Fig. 3d). Additionally, SCARLink predicted numerous false-positive CRRs in monocytes, further emphasizing its reduced specificity relative to SCRIPT (Fig. 3d). Notably, SCRIPT successfully identifies a B-cell-specific CRR spanning 16.5 Mb upstream of the *CCR6* TSS, a regulatory interaction missed by both SCARLink and LINGER (Fig. 3e, Fig. S7b).

Similar trends are observed at other gene loci, such as *PIK3R5, EFNA5, ENPP2, CDCA7L, IRAK3, SYTL2*, and *AOAH* (Fig. S8-11). Taken together, these results underline SCRIPT’s superior capacity to identify long-range CRRs with high precision and specificity, outperforming both SCARLink and LINGER.

### SCRIPT attends to cell-type-specific enhancers

To determine whether CREs identified by SCRIPT are preferentially regarded as enhancers, we examine the overlap between enhancers labeled by H3K27ac ChIP–seq data and CRRs predicted by SCRIPT. We find that those CRRs with astrocyte-specific enhancers exhibit significantly higher regulation scores in astrocytes compared to the other cell types (Fig. S12a). Similar trends are observed for enhancers specific to microglia, neurons, and oligodendrocytes (Fig. S12b-d). In addition, we identify marker CRRs and marker genes for each cell type through differential analysis of the regulation score matrix and the gene expression matrix. The analysis shows a significant concordance between genes regulated by marker CRRs and marker genes for the same cell type (Fig. S12e). These results suggest that CRRs with high regulation scores are likely to promote the expression of their corresponding target genes, strongly indicating that SCRIPT prioritizes the identification of cell-type-specific enhancer-like CREs. Similar patterns are observed in the PBMC dataset (Fig. S13).

### SCRIPT understands the pathogenic mechanisms of AD and SCZ in a cell-type-specific manner

Both AD and SCZ are complex genetic diseases characterized by progressive cognitive impairment and structural alteration in the brain. GWASs conducted by Iris E. Jansen et al.^25^ and Antonio F. Pardiñas et al.^26^ have identified 275 AD-associated and 1,895 SCZ-associated single nucleotide polymorphisms (SNPs) mapped to CREs identified through scATAC-seq data in the human brain^27^ (Table S3-4). Notably, the majority of these SNPs are located in non-coding regions of the human genome.

SCRIPT offers a novel approach to explain the pathogenic mechanisms of non-coding SNPs identified by GWAS. By linking SNPs to their target genes at the single-cell resolution, SCRIPT identifies cell types contributing to genetic risk from non-coding SNPs (Table S5-6). It further elucidates the biological functions influenced by the SNPs and nominates the target genes of the SNPs specific to the cell types of interest (Fig. 4a).

**Fig. 4.**
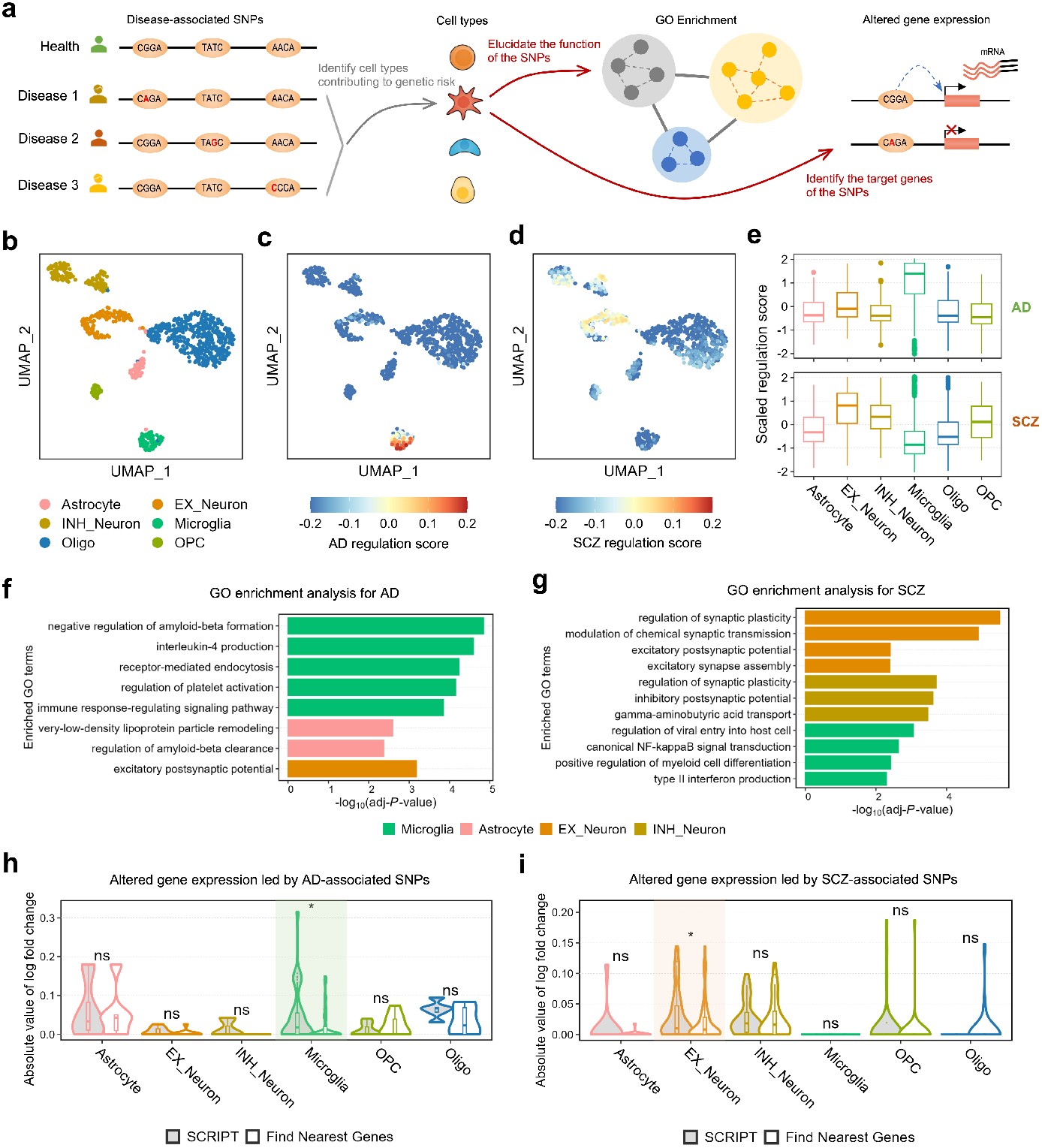
SCRIPT elucidates the pathogenic mechanisms of AD-and SCZ-associated variants. **a,** Schematic representation of the overall strategy for investigating the pathogenic mechanisms of disease-associated variants at the cell-type level using SCRIPT. **b,** UMAP visualization of scATAC-seq data from the human cortex dataset including six cell types (*n* = 852 cells). **c, d,** UMAP visualization of single-cell mean regulation scores for CRRs related to AD **(c)** and SCZ **(d). e,** Boxplots of scaled regulation scores of each human cortex cell type at the CRRs involved in AD- and SCZ-related variants (AD: 141 CRRs; SCZ: 392 CRRs). **f, g,** Bar plots displaying the enrichments of GO terms in genes regulated by AD- and SCZ-related variants across different cell types. **h, i,** Violin plots illustrating the absolute values of log fold changes (ALFCs) in gene expression between disease states (AD **(h)** or SCZ **(i)**) and controls. Genes identified by SCRIPT and genes determined using the nearest-gene approach are represented in gray and white, respectively, within the violin plots. One-sided Wilcoxon rank sum tests are performed between ALFCs of genes identified by SCRIPT and those of genes determined using the nearest-gene approach (\**P* < 0.05). SNP, single nucleotide polymorphism; GO, gene ontology; AD, Alzheimer’s disease; SCZ, schizophrenia.

The regulation score of each single cell is calculated by averaging the regulation scores of CRRs related to AD or SCZ per cell. Cells labeled as microglia exhibit the highest regulation scores for AD, while excitatory neurons show the highest regulation scores for SCZ (Fig. 4b-d). Besides, the regulation score for each CRR was determined by averaging the regulation scores of cells of the same type for CRRs related to AD or SCZ. Consistently, the highest CRR regulation scores were observed in microglia for AD and in excitatory neurons for SCZ (Fig. 4e). These results reflect that microglia and excitatory neurons are the primary cell types contributing to genetic risk for AD and SCZ, respectively, which is supported by the literature of AD and SCZ^28, 29^.

Subsequently, we select the genes regulated by the cell-type-specific CRRs related to SCZ or AD to perform gene ontology (GO) enrichment analysis. For AD, Genes specifically regulated in microglia are enriched in gene ontology (GO) terms such as “negative regulation of amyloid-beta formation”, “receptor-mediated endocytosis”, and “regulation of platelet activation”, while astrocyte-specific regulated genes are enriched in “very-low-density lipoprotein particle remodeling” and “regulation of amyloid-beta clearance” (Fig. 4f). Extracellular β-amyloid (Aβ) deposition is a pathological hallmark of AD^30^. Activated microglia encase Aβ plaques to form a barrier that limits dissemination and can phagocytose Aβ, thereby attenuating Aβ-driven neuronal damage. These cellular defense processes have crowned microglia as perhaps the most important cell type in the control of amyloid burden in AD^28^. Besides, platelet activation increases plasma Aβ levels, contributing to AD pathogenesis^31^. Additionally, Aβ and astrocytes-derived APOE form an APOE-Aβ complex which is cleared by a very-low-density lipoprotein receptor at the blood-brain barrier^32^.

For SCZ, the results of GO enrichment analysis indicate that the excitatory-neuron-specific regulated genes are enriched for “regulation of synaptic plasticity”, “modulation of chemical synaptic transmission” and so on. The inhibitory neuron-specific regulated genes are enriched for “regulation of synaptic plasticity”, “gamma-aminobutyric acid transport” and so on. Microglia-specific regulated genes are enriched for “canonical NF-kappaB signal transduction”, “type II interferon production” and so on (Fig. 4g). Synaptic plasticity and neurotransmitter transmission have been suggested to be impaired in SCZ^33, 34^, and neuroinflammation is considered to contribute to the pathogenesis of SCZ^35^. Overall, these pathway enrichment results are consistent with the literature on AD and SCZ.

Finally, scRNA-seq data for AD and SCZ populations^29, 36^ demonstrate that SCRIPT identifies more biologically relevant target genes for AD- and SCZ-associated SNPs at the cell-type level. In microglia, the AD-related genes identified by SCRIPT exhibit significantly greater differential expression than the nearest genes to AD-associated SNPs (Fig. 4h). Similarly, in excitatory neurons, the SCZ-related genes identified by SCRIPT show significantly greater differential expression than the genes closest to SCZ-related SNPs (Fig. 4i).

### SCRIPT elucidates the function of disease-related SNPs by nominating target genes

We present some analytical cases using SCRIPT to elucidate the function of AD-or SCZ-associated SNPs by nominating their target genes. SCRIPT predicts nine microglia-specific CRRs between *APOE* and nine CREs harboring 19 AD-associated GWAS SNPs. These CREs are located downstream of the *APOE* TSS, spanning a range from 6 kb to 174 kb. One illustrative example is rs7251911, a SNP located in the intron of *GEMIN7*. SCRIPT predicts that this SNP regulates *APOE*, located 174 kb upstream, through a microglia-specific CRR (Fig. 5a, Fig. S14). *APOE* exhibits high expression levels in microglia and astrocyte (Fig. 5b,c). Besides, *APOE* remains the most significant genetic risk factor for AD, implicated in processes such as Aβ peptide aggregation and clearance, tau neurofibrillary degeneration, microglial and astrocytic responses, and blood-brain barrier disruption—all of which contribute to cognitive decline^32^. In contrast, the transcriptional levels of *GEMIN7* are minimal across cortical cell types (Fig. 5b,d). In this case, SCRIPT identifies a more plausible mechanism for AD-causing genetic variants by assigning rs7251911 to *APOE*, rather than *GEMIN7*. Notably, PLAC-seq data also link rs7251911 to *APOE*, whereas other methods like LINGER and SCARLink fail to identify this long-range CRR (Fig. S15).

**Fig. 5.**
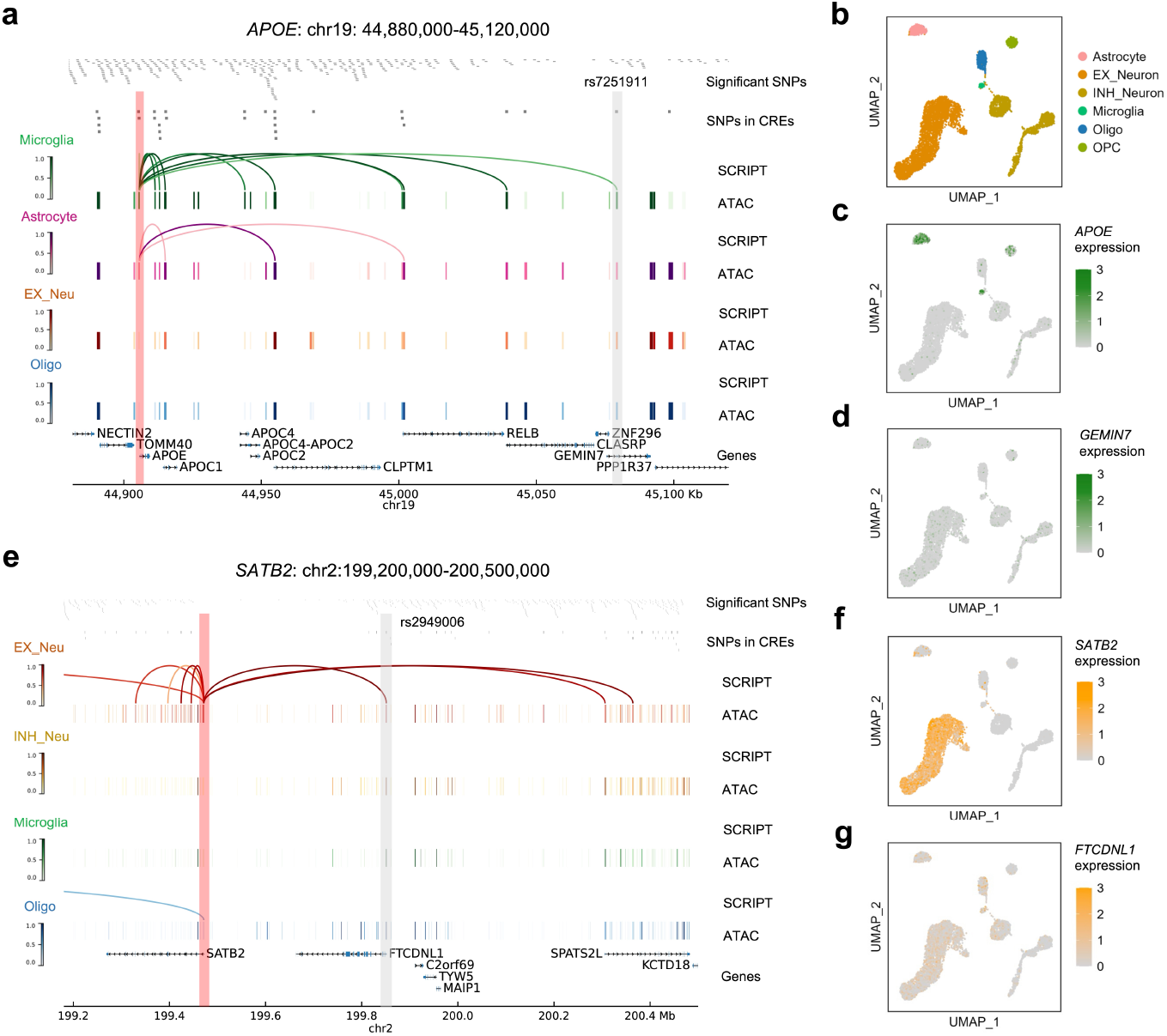
Application of SCRIPT to nominate gene target for AD- and SCZ-related SNPs. **a,** Genomic visualization of normalized regulation scores predicted by SCRIPT and normalized scATAC– seq-derived pseudobulk tracks across four cell types in the *APOE* gene locus (chr19:44,880,000-45,120,000). SNPs significantly associated with AD and located within CREs are displayed at the top of this panel. The genomic locations of SNPs of interest and the *APOE* transcription start site (TSS) are highlighted by gray and red bars, respectively. **b,** UMAP visualization of scRNA-seq data from the human cortex dataset including six cell types (*n* = 15,594 cells). **c, d,** UMAP visualization of *APOE* **(c)** and *GEMIN7* **(d)** expression across six cortex cell types. **e,** Genomic visualization of the *SATB2* gene locus (chr2:199,200,000–200,500,000) and SCZ-related SNPs, following the same organization as **Fig. 5a. f, g**, UMAP visualization of *SATB2* **(f)** and *FTCDNL1* **(g)** expression across six cortex cell types.

In another analytical case, three CREs containing seven SCZ-associated GWAS SNPs, located downstream from 379 Kb to 894 Kb, are predicted to regulate *SATB2* through excitatory-neuron-specific CRRs. A representative example is rs2949006, an SNP located in the promoter of *FTCDNL1*. This SNP is predicted by SCRIPT to regulate *SATB2* through an excitatory-neuron-specific CRR (Fig. 5e, Fig. S16). *SATB2* is associated with SCZ risk and is an important transcription factor regulating neocortical organization and circuitry^37^. In contrast, *FTCDNL1* exhibits minimal transcriptional levels across cortical cell types (Fig. 5f, g). By linking rs2949006 to *SATB2*, SCRIPT identifies SATB2 as a more likely SCZ-causing gene instead of *FTCDNL1*. PLAC-seq data of neurons confirm this assignment, while LINGER and SCARLink fail to detect the long-range CRR (Fig. S17).

Additionally, we employ SCRIPT to identify more cases in *BIN1, RIN3, MS4A6A, MS4A7, HCN1, MMP16, NRGN*, and *SLC32A1* loci (Fig. S18-21). Previous studies have supported the associations between these genes and AD or SCZ^38–45^.

These cases demonstrate that SCRIPT provides a more comprehensive prioritization of putative functional SNPs at the cell-type level compared to traditional approaches, such as LINGER, SCARLink, and nearest-gene assignment (Fig. 1e, middle). By prioritizing functional SNPs and nominating their target genes, SCRIPT enhances our understanding of the functions of non-coding SNPs. For instance, SNPs located in distinct CREs may regulate the same target gene through long-range *cis*-regulation, as predicted by SCRIPT, suggesting potential shared molecular genetic mechanisms. In contrast, traditional methods may overlook these connections, potentially obscuring insights into patient stratification of complex genetic diseases (Fig. 1e, right).

## Discussion

SCRIPT is a method that leverages GCAT underpinned by comprehensive empirical CRR evidence, combined with representation learning driven by large-scale scATAC-seq data. First, SCRIPT employs GCAT to simulate *cis*-transcriptional regulation, offering a biologically interpretable alternative to traditional black-box deep learning models. GCATs selectively update the attention scores of CRE-gene links supported by empirical CRR evidence. Therefore, the simulation bridges scATAC-seq and scRNA-seq data guided by causal biological principles, which prevents SCRIPT from the noises introduced by data correlation. GCAT ensures the reliability of SCRIPT, which not only minimizes false positives but also effectively captures long-range CRRs. Second, SSGAE enhances CRE representations by effectively capturing the complex interactions within large-scale scATAC-seq data. These enhanced representations contribute to improved accuracy in gene expression predictions, further strengthening SCRIPT’s ability to identify cell-type-specific CRRs. These innovative algorithmic designs ensure that SCRIPT achieves substantially superior CRR inference performance, especially for long-range CRRs.

More importantly, we present a systematic framework to prioritize candidate disease-causing SNPs from numerous non-coding GWAS SNPs at cell type level and elucidate the functions of the disease-causing SNPs by nominating with high confidence gene and cellular targets for them. Compared to current computational methods, such as LINGER and SCARLink, SCRIPT demonstrates superior accuracy in identifying long-range CRRs, enabling a more comprehensive annotation of the functional effects of non-coding SNPs. For example, SCRIPT assigns the AD-related SNP rs7251911 to *APOE* specific in microglia, and the SCZ-related SNP rs2949006 to *SATB2* specific in excitatory neurons. In contrast, conventional approaches, which cannot detect long-range CRRs and rely solely on nearest-gene assignment in a cell-type-agnostic manner, would misattribute these SNPs, limiting their utility for precise genetic-based patient stratification and novel therapeutic target discovery. More broadly, this study establishes an effective approach for identifying individuals at genetic risk by prioritizing more disease-causing non-coding SNPs with high specificity. Furthermore, it provides an avenue towards the nomination of new therapeutic targets that previously remained obscured by the cell heterogeneity and complexity of the regulatory machinery of the noncoding genome.

While SCRIPT offers significant advantages, it also has two limitations that merit closer consideration. First, while SCRIPT incorporates comprehensive empirical CRR evidence as causal biological principles, its reliance on experimentally measured CRRs limits its ability to predict CRRs beyond current experiments. Expanding the scale of the empirical CRR evidence through broader data collection and the development of advanced bulk CRR prediction algorithms will address this limitation. Second, SCRIPT only employs scATAC-seq data for single-cell CRR prediction.

Incorporating additional single-cell epigenomic datasets, such as methylation and histone modification data, could provide deeper insights into gene regulation and enhance the overall utility of the framework.

## Materials and Methods

### The inputs for SCRIPT

#### Input data

SCRIPT requires two input matrices: a well-annotated scATAC-seq count matrix and a corresponding scRNA-seq count matrix. Notably, the two matrices do not need to originate from single-cell multi-omic data where both scRNA-seq and scATAC-seq are applied to the same individual cells, but they must be derived from the same tissue. Additionally, SCRIPT relies on a comprehensive empirical CRR evidence database, constructed through the integration of large-scale bulk Hi-C and eQTL data.

#### Preprocessing

In this study, the reference genome version for the scATAC-seq matrix is hg38/GRCh38. For datasets based on other reference genome versions, conversion to hg38/GRCh38 can be performed using LiftOver.

Quality control is applied to the scATAC-seq matrix. By default, cells with fewer captured peaks than the 1st percentile or more than the 99th percentile are excluded. At the gene level, peaks with less than 5% occurrence within any cell type are removed.

Because the low sequencing depth of scATAC-seq data may hinder CRR predictions, data aggregation is performed. In detail, the scATAC-seq count matrix is normalized, and 50,000 highly variable features are selected. After scaling the data, latent semantic indexing (LSI) is used to reduce the data to 50 dimensions. These dimensions are then employed to identify the 10 nearest neighbors for each cell, and the chromatin accessibility profiles of the 11 cells (the target cell and its 10 neighbors) are aggregated.

The scRNA-seq count matrix is transformed into gene expression profiles of each cell type by aggregating the expression profiles of single cells with the same cell label. For the gene expression profiles, cosg^46^ is used to identify 300 marker protein-coding genes for each cell type, generating a marker gene list. Only genes within this list are considered for subsequent modeling. To mitigate the influence of library size, the expression profile matrix is normalized by cell type such that the sum of all gene expression values within each cell type equals 1.

#### Construction of cell graphs

The edges of cell graphs are derived from a comprehensive empirical CRR evidence database, which integrates data from three sources: proximal CRRs, distal CRRs from pcHi-C data, and CRRs from eQTL data. Proximal CRRs are defined as the CRRs between a gene and the CREs located within its promoter region. The promoter region is specified as the genomic interval spanning 2 kb upstream to 2 kb downstream of TSS. A CRE is considered to be located within the promoter region if it overlaps with the region by at least one bp. CRRs from pcHi-C data are generated from interactions between promoter fragments and either other promoter fragments or distal fragments. Interactions between promoter fragments produce CRRs linking genes and CREs within promoter regions, while interactions between promoter fragments and distal fragments generate CRRs linking genes and CREs in distal fragments. A CRE is considered to belong to a promoter or distal fragment if it overlaps by at least one bp. CRRs from eQTL data are based on associations between genes and SNPs. A CRE is considered to regulate a gene if it overlaps by at least one bp with an SNP linked to the gene. All positional intersection operations are implemented by bedtools^47^.

In the pretraining stage, cell graphs consist solely of CRE nodes. For each gene, all CREs within its promoter region are connected to CREs linked to the gene in the CRR database. In the cis-regultion simulation stage, nodes are categorized into genes and CREs, and edges between these nodes are established based on the CRR database. For a given cell graph, the values of CRE nodes are derived from the chromatin accessibility vectors of the corresponding cell in the scATAC-seq matrix. The values of gene nodes are determined based on the chromatin accessibility vectors of CREs within the promoter region closest to the TSSs of the respective genes.

### The pretraining of self-supervised graph autoencoder

During the pretraining stage, scATAC-seq data consisting of *N* cells that can be converted into *N* cell graphs. We use 𝒢*^p^* to denote the set of cell graphs used in the pretraining stage and 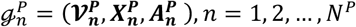 to denote the *n*th cell graph where 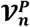 represents the node set,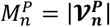 is the number of nodes, 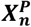 represents chromatin accessibility vector and 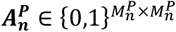 represents the adjacency matrix of CRE-CRE links. Further, given *f_E_* as the graph encoder, *f_D_* as the graph decoder, and 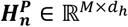 denoting the hidden state of the *n*th cell graph encoded by the encoder, the goal of general self-supervised graph autoencoder (SSGAE) is to reconstruct the input as

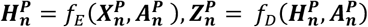

where 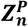 denotes the reconstructed features.^16^

In SCRIPT, GAT^14^ is used as the backbones of *f_E_* and *f_D_* to propagate information between nodes in each cell graph. Given a cell graph, GAT takes as input the hidden state matrix 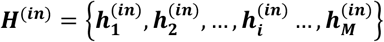, where 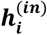 is the hidden state vector of the *i*th node. The updated hidden state matrix ***H***^(***out***)^ is computed as:

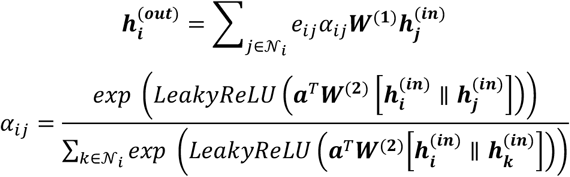

Here, 𝒩*_i_* is the set of the neighboring nodes of the *i*th node in the graph 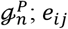 is the weight of the edge linking the *i*th node and the *j*th node; *α_ij_* is the attention score between the *i*th node and the *j*th node; ***W***^(**1**)^ and ***W***^(**2**)^ are learnable parameter matrices; ***a*** is a learnable parameter vector.

In the masking stage of SSGAE, for a given cell graph 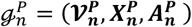, a subset of nodes 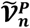 is randomly selected from 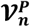, and their features are replaced with a mask token ***x***_[***M***]_. The masked node feature 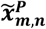 for 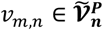 in the masked feature matrix 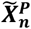 is defined as:

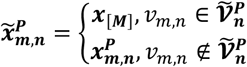

The objective of SSGAE is to reconstruct the masked nodes features in 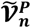 given the partially observed node signals 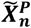 and the input adjacency matrix 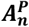.

In the reconstruction stage of SSGAE, scaled cosine error (SCE) is leveraged as the loss function to reconstruct original node features. Given the original feature 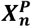 and reconstructed output 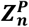, the SCE for SSGAE is defined as:

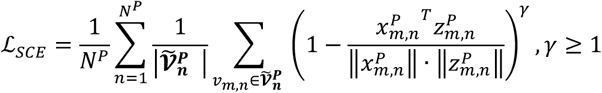

where 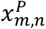 and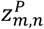 respectively represent the components of 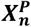 and 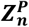 in the *m*-th node *v_m,n_* of *n*-th cell, and the easy samples’ contribution is down-weighted during training by scaling the cosine error with a power of *γ* ≥ 1.

### *Cis*-regulation simulation using graph causal attention networks

In a specific tissue, a matched scATAC-seq and scRNA-seq dataset (with aligned cell types) is leveraged to train a *cis*-regulation simulation model. Each cell of scATAC-seq data is transformed into a cell graph 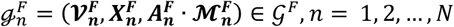, where 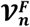 represents the node set, 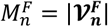 is the number of nodes, 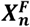 represents chromatin accessibility vector, 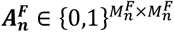 represents the adjacency matrix of CRE-gene links, and 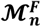 represents the causal attention mask matrix^15^. Each element 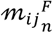 in the *i-*th row and *j*-th column of 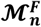 is assigned a value of 0 or

1 according to the following rule:

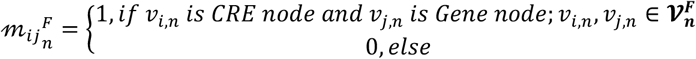

For each cell graph 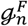, a linear layer followed by a RELU activation function encodes 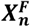 into the hidden state matrix 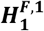, while SSGAE to encode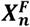 to the hidden state matrix 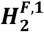. The GCAT layer *φ* is employed to obtain the updated node hidden state matrix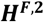as follows:

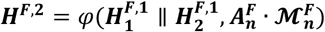

Then, ***H*^*F*,2^** is pooled into a vector 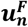 with the same size as 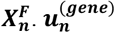 is obtained by selecting the gene nodes from 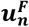. Then, 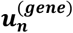 is normalized as follows:

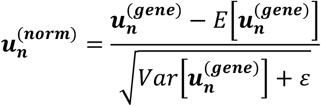

where *E*[·] is the calculation of mathematical expectation, *Var* [·] is the calculation of variance, *ε* is a value added to the denominator for numerical stability. Here, 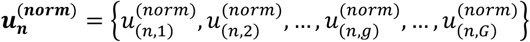 contains *G* values, and 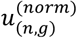 is the normalized predicted value of the *g*th gene in the *n* th cell.

Subsequently, a softmax function is used to transform each element of 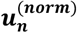 into a probability as follows:

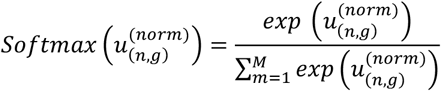

This transformation yields 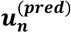, representing the predicted probability vector of sequence reads in each gene region. The true probability vector of the *n*th cell is denoted as 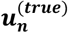. The first part of the loss function of the *cis*-regulation simulation model, termed the gene expression loss function, is calculated as follows:

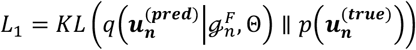

where *KL*(*q*(·) ∥*p*(·))is the Kullback–Leibler divergence between predicted distribution 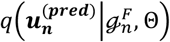 and prior distribution 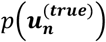, and Θ is the parameters of the *cis*-regulation simulation model.

Additionally, 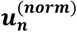 serves as input to a multilayer perceptron to generate the label probability vector of the *n-*th cell ***y_n_***={*y*_(*n*,1)_,*y*_(*n*,2)_, …*y*_(*n*,*c*)_, …*y*_(*n*,*C*)_}, where the model predicts one of seven possible labels and *y*_(*n*,*c*)_ represents the probability that the *n*th cell belongs to the *c-*th label. The true cell-type label vector is denoted as **𝒞*_n_*** = {𝒞_(*n*,1)_, 𝒞_(*n*,1)_, … 𝒞_(*n*,2)_, … 𝒞_(=*n*,*C*)_}, where 𝒞_(*n*,*c*)_ is 1 if the *n-*th cell belongs to class *c*, and 0 otherwise. The second part of the loss function of the *cis*-regulation simulation model, the label loss function, is defined as the weighted cross-entropy between the predicted and true label vector:

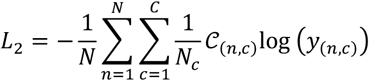

where 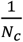 is the weight of the *c-*th label, *Nc* is the number of the *c-*th label in the training dataset.

Finally, the total loss function is defined as:

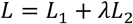

where *λ* balances the contributions of *L*_1_ and *L*_2_ in model optimization and is set to 1 by default.

### The regulation score prediction based on the attribution method

In the test dataset, we use the well-trained *cis*-regulation simulation model to predict the gene expression of each cell. To infer single-cell regulation scores, we apply an attribution method, integrated gradients (IG)^48^, which quantifies the influence of changes of edge weights in each cell graph on the expression levels of regulated genes.

For the *g*th gene in the *n*th cell, 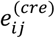 represents an edge weight in the CRR networks *e*^(*cre*)^. The influence of a change of 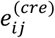 on the expression level of the *g*th gene is calculated as:

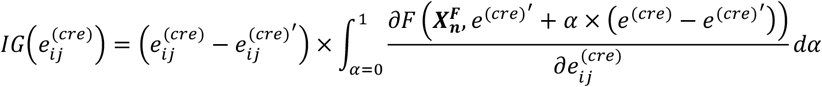

where *F*(·) denotes the predicted expression level of the *g*th gene, and 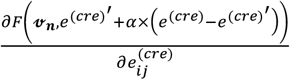 is the gradient of *F*(·) with respect to 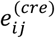, computed when the node feature vector is 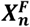 and the edge weights of the cell graph is 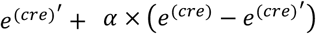. By default, we set all elements of *e*^(*cre*)^ as 1 and set all elements of baseline 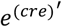 as 0. The calculation of attribution scores can be simplified as:

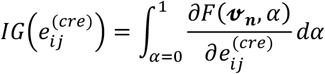

Using this formula, we compute the attribution scores for all edges connected to the *g*th gene in the *n*-th cell. This process is repeated for each gene in the *n*-th cell, yielding an attribution score vector with a length equal to the number of edges in the cell graph.

Finally, we compute the regulation score (RS) of each CRR per cell by normalizing the corresponding attribution score to a range between −1 and 1, using the following transformation:

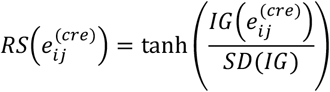

where tanh is a hyperbolic tangent function, and *SD*(*IG*) is the standard deviation of all attribution scores.

## Data collections

All datasets used in this study are downloaded from public databases.

The construction of the empirical CRR evidence database relies on large-scale pcHi-C and eQTL datasets, sourced as follows:

- The large-scale pcHi-C dataset is obtained from the study by Inkyung Jung et al^4^, with accession number GSE86189 in the GEO database. In the study, the pcHi-C technique was used to obtain chromatin interaction data in the 27 human cell/tissue types.
- The *cis*-eQTL dataset, derived from 49 human tissues, is provided by the GTEx project^10^ and is available at https://www.gtexportal.org/.

The human scATAC-seq atlas from the study by Kai Zhang et al^49^ is utilized to pretrain the SSGAE, and its accession number in the GEO database is GSE184462.

Several scRNA-seq and scATAC-seq datasets are used to predict single-cell CRRs, and their sources are as follows:

- The scRNA-seq dataset of the middle temporal gyrus of the human cortex is obtained from the study by Rebecca D. Hodge et al^50^and is available at http://celltypes.brain-map.org/.
- The scATAC-seq dataset of the human cortex is obtained from the study by Ryan M. Mulqueen et al^27^ (GEO accession number: GSE174226).
- The scRNA-seq and scATAC-seq datasets of the human prefrontal cortex are from the study by Samuel Morabito et al^51^, with accession number GSE174367 in the GEO database.
- The scRNA-seq and scATAC-seq datasets of human peripheral blood mononuclear cells (PBMC) are sourced from 10X Genomics website and are downloaded from https://support.10xgenomics.com/single-cell-gene-expression/datasets.
- The scRNA-seq and scATAC-seq datasets of human bone marrow mononuclear cells (BMMC) and PBMC are derived from the study by Jeffrey M. Granja et al^52^, and their accession number in the GEO database is GSE139369.

To validate the predicted cell-type-specific CRRs, two cell-type-specific chromatin contact datasets and two cell-type-specific ChIP-seq datasets were utilized, sourced as follows:

- The PLAC-seq and H3K27ac ChIP-seq data for three human cortex cell types are obtained from the study by Alexi Nott et al^19^ and are available in supplementary table S5 of their paper.
- The pcHi-C data for 17 human blood cell types are obtained from the study by Biola M. Javierre et al^20^ and are available in supplementary table S1 of their paper.
- The H3K27ac ChIP-seq data of four immune cell types are obtained from the Roadmap Epigenomics Project^24^ and are available at https://egg2.wustl.edu/roadmap/web_portal/processed_data.html.

Two GWAS summary statistics datasets are employed to explore the pathogenic mechanisms of AD and SCZ, with sources as follows:

- GWAS summary statistics data for AD are obtained from meta-analysis by E. Jansen et al.^25^ (71,880 cases, 383,378 controls) and are available at https://ctg.cncr.nl/software/summary_statistics.
- GWAS summary statistics data of SCZ are obtained from meta-analysis by Antonio F. Pardiñas et al.^26^ (40,675 cases, 64,643 controls) and are available at https://ctg.cncr.nl/software/summary_statistics.

The scRNA-seq data of AD and SCZ populations are sourced from the studies by Hansruedi Mathys et al.^36^ and W. Brad Ruzicka et al.^29^, respectively. Differential expression gene lists across different cell types for AD and SCZ are extracted from supplementary tables of the two studies.

### Implementation details

For SSGAE, the encoder consists of two GAT layers, while the decoder contains one GAT layer. The *cis*-regulation simulation model is implemented with a single GAT layer. In all GAT layers used in this study, the node hidden state has a dimension of 64, the number of multi-head attentions is set to 4, and the dropout probability for the normalized attention coefficients—allowing each node to be exposed to a stochastically sampled neighborhood during training—is set to 0.5. The masking ratio for SSGAE is also set to 0.5.

Model parameters are optimized using the Adam algorithm, with initial learning rates of 1 × 10^−4^ for SSGAE and 5 × 10^−4^ for the *cis*-regulation simulation model. To prevent overfitting, L2 regularization is applied with a weight decay parameter of 1 × 10^−4^.

During the *cis*-regulation simulation, for each scATAC-seq dataset, we randomly select 50% of the cells as the training set and 10% as the validation set to train the gene expression model. The well-trained model is then used to predict the gene expression matrix for the remaining 40% of test cells. Finally, the attribution model is applied to the test set to obtain regulatory scores.

### Evaluation for cell-type-specific CRR prediction

To assess the accuracy of cell-type-specific CRR predictions, we employ two metrics: cell-level AUC and reg-level AUC.

AUC quantifies a classifier’s ability to distinguish between two classes. It is computed by plotting the true positive rate (TPR) against the false positive rate (FPR) across different thresholds. The AUC value represents the probability that a randomly selected positive example is ranked higher than a randomly selected negative example. A perfect classifier has an AUC of 1, while a random classifier has an AUC of 0.5.

The result of CRR prediction is a regulation score matrix, where rows correspond to cell types and columns to CRRs. Cell-level AUC evaluates whether the regulation scores of a given cell type can effectively distinguish cell-type-specific CRRs from non-specific CRRs. Reg-level AUC assesses whether the regulation scores of an experimentally validated cell-type-specific CRR can distinguish the corresponding cell type from others.

For comparisons between predicted CRRs and cell-type-specific chromatin contact data, we merge cell subtypes to ensure consistency in cell-type annotations. In brain tissue, excitatory and inhibitory neurons are combined into a single neuron category. In the PBMC tissue, we consolidate subtypes as follows: CD4^+^ naïve T cells, CD4^+^ central memory T cells, CD4^+^ effector memory T cells, and regulatory T cells are grouped into CD4^+^ T cells; CD8^+^ naïve T cells and CD8^+^ effector memory T cells are merged into CD8^+^ T cells; CD14^+^ monocytes and CD16^+^ monocytes are combined into monocytes; and naïve B cells, intermediate B cells, and memory B cells are categorized as B cells.

### Identification of marker CRRs and marker genes

We identify marker CRRs for each cell type as follows. For each CRR in the single-cell regulation score matrix, we first compute the intra-group distances of regulation scores within a given cell type, followed by the inter-group distances of regulation scores between this cell type and all others. The fold change (FC) value is defined as the mean inter-group distances divided by the mean intra-group distance. To assess statistical significance, we apply the Wilcoxon rank-sum test to determine whether the inter-group distances are significantly greater than the intra-group distances. A CRR is deemed specific to a cell type if the FC value exceeds 1.5 and the Bonferroni-corrected *P*-value is below 0.05.

We perform the FindAllMarkers function of the Seurat package^53^ to identify marker genes of each cell type from a single-cell gene expression matrix. Only the genes with FC value >1.5 and Bonferroni-corrected *P*-value < 0.05 were regarded as marker genes.

### Visualization of CRR prediction

For the predicted single cell regulation score matrix, we select 3000 highly variable CRRs, and performance PCA to reduce this matrix to 50 dimensions. We then apply UMAP for the 50 PCs to visualize the single-cell regulation score matrix to compare the performance of three methods in capturing biologically relevant signals related to cell type.

To visualize CRRs and scATAC-seq peaks in specific genomic regions, we use pyGenomeTracks^54^. Only predicted CRRs with regulation scores greater than 0.05 are displayed. For each visualized gene locus, the regulation scores generated by each computational method are normalized by dividing by their respective maximum values. For the cell-peak matrix derived from scATAC-seq data, we generate pseudobulk scATAC-seq profiles by aggregating single cells belonging to the same cell type. The pseudobulk scATAC-seq data are then normalized using a size factor of 10^5^. Since most normalized accessibility scores fall within the range of 0 to 1, values exceeding 1 are clipped to 1.

### Application of SCRIPT to explain disease-associated variants

We define variants with *P* values less than 1 × 10^−5^ in GWAS as disease-associated SNPs. To analyze these variants, we select scATAC-seq and scRNA-sq data from disease-relevant tissues and apply SCRIPT to generate a single-cell regulation score matrix for the tissue. Using bedtools, we map disease-associated SNPs to CREs identified in the scATAC-seq data. We then extract CRRs involving these disease-associated CREs to construct a disease-associated single-cell regulation score matrix. The regulation score vector for each cell type is obtained by averaging the regulation score vectors of all cells belonging to that cell type. To facilitate comparison across CRRs, we perform a Z-transformation on the regulation scores of all cell types for each CRR, yielding scaled regulation scores.

To elucidate the biological functions of disease-associated variants, we identify genes regulated by cell-type-specific CRRs and conduct GO enrichment analysis. GO enrichment analysis is performed using ClusterProfiler^55^, with multiple testing corrections applied via the Benjamini-Hochberg method at a significance threshold of 0.05.

To evaluate whether the target genes identified by SCRIPT are more biologically relevant to AD or SCZ, we analyze scRNA-seq data from AD and SCZ cohorts obtained from studies conducted by Hansruedi Mathys et al.^36^ and W. Brad Ruzicka et al.^29^, respectively. These studies provide the results from differential gene expression analysis (DGEA) for each cell type, comparing control samples with either AD or SCZ samples. For disease-associated variants of each disease, two target gene lists are generated: one using SCRIPT and the other by identifying the nearest gene to each variant. The absolute values of log fold changes for both gene lists are extracted from the DGEA results. To determine whether genes identified by SCRIPT exhibit significantly higher absolute log fold changes compared to those from the nearest-gene approach, we apply the Wilcoxon rank-sum test. Notably, the absolute log fold change of a gene is set to 0 if its false discovery rate (FDR) exceeds 0.05.

